# Molecular species delimitation in the primitively segmented spider genus *Heptathela* endemic to Japanese islands

**DOI:** 10.1101/812214

**Authors:** Xin Xu, Matjaž Kuntner, Jason E. Bond, Hirotsugu Ono, Simon Y. W. Ho, Fengxiang Liu, Long Yu, Daiqin Li

## Abstract

Determining species boundaries forms an important foundation for biological research. However, the results of molecular species delimitation can vary with the data sets and methods that are used. Here we use a two-step approach to delimit species in the genus *Heptathela*, a group of primitively segmented trapdoor spiders that are endemic to Japanese islands. Morphological evidence suggests the existence of 19 species in the genus. We tested this initial species hypothesis by using six molecular species-delimitation methods to analyse 180 mitochondrial *COI* sequences of *Heptathela* sampled from across the known range of the genus. We then conducted a set of more focused analyses by sampling additional genetic markers from the subset of taxa that were inconsistently delimited by the single-locus analyses of mitochondrial DNA. Multilocus species delimitation was performed using two Bayesian approaches based on the multispecies coalescent. Our approach identified 20 putative species among the 180 sampled individuals of *Heptathela*. We suggest that our two-step approach provides an efficient strategy for delimiting species while minimizing costs and computational time.

## 1. Introduction

Species delimitation is an important process that underlies a wide range of biological questions (Agnarsson and Kuntner, 2007; Camargo and Sites, 2013). However, conflicts in species delimitation can be caused by differences in the application of species concepts (de Queiroz, 2007; Freudenstein et al., 2017) and in the choices of methods, models, and genetic markers (Carstens et al., 2013; Schlick-Steiner et al., 2010, 2014). Furthermore, there are differing views on the existence and importance of cryptic species (Fišer et al., 2018; Heethoff et al., 2018; Struck et al., 2018). This can pose challenges for the study of taxa that are well differentiated genetically but not morphologically (Bond et al., 2001; Leavitt et al., 2015). Resolving these conflicts is crucial for the advancement of integrative taxonomy.

Taxa inhabiting islands, mountains, and caves represent cases where lineages might be morphologically homogeneous but highly genetically structured. Island populations can experience morphological stasis as a result of shared selective pressure or niche conservatism (Gillespie and Roderick, 2002; Opatova and Arnedo, 2014). However, barriers to gene flow among isolated islands and among habitats within islands can lead to deep genetic structuring (Gillespie and Roderick, 2002; Opatova and Arnedo, 2014; Xu et al., 2016). The combination of morphological homogeneity with population genetic structuring can be seen in taxa with limited dispersal capability and high endemism (Derkarabetian et al., 2019; Hedin et al., 2015), thus presenting special challenges for species delimitation. Such conditions are seen in the Japanese archipelagos, which are among the most threatened biodiversity hotspots in the world (Crowe et al., 2006; Myers et al., 2000). These islands harbour diverse and largely endemic floras and faunas, including the primitively segmented trapdoor spiders (Araneae: Liphistiidae).

Liphistiidae is the only extant family in the suborder Mesothelae, an ancient and species-poor lineage of spiders that contains species with disjunct geographical distributions restricted to Southeast and East Asia (World Spider Catalog, 2020; Xu et al., 2015a, 2015b). The family has eight genera, each endemic to a particular geographical region. Like other arachnids (Derkarabetian and Hedin, 2014) and other trapdoor spiders (Bond et al., 2001; Satler et al., 2013), liphistiids often show limited dispersal ability and high habitat specificity (Haupt, 2003; Xu et al., 2015a). The biogeographic patterns in liphistiids make them an ideal system for studying speciation. However, species of liphistiids have historically been difficult to delimit because of their uniform, conservative morphology.

Liphistiid systematics and taxonomy are in need of revision, primarily due to insights from a recent series of molecular phylogenetic studies (Xu et al., 2015a, 2015b, 2015c, 2016, 2017). The efficacy of single-locus analyses based on the mitochondrial *COI* barcode region has been confirmed repeatedly by delimitation of species in the liphistiid genera *Ganthela* Xu & Kuntner, 2015 and *Ryuthela* Haupt, 1983, enabling nomenclatural changes to be made (Xu et al., 2015c, 2017). Multilocus methods for species delimitation are expected to provide a more reliable identification of species boundaries, because they can collect information from multiple independently evolving loci. However, some studies have found that such methods detect population structure rather than species divergences, thus leading to oversplitting of species diversity in short-range endemic taxa (Chambers and Hillis, 2020; Hedin et al., 2015). In spite of this potential weakness, multilocus approaches can provide valuable information for guiding taxonomic decisions.

Here we focus on the liphistiid spider genus *Heptathela* Kishida, 1923 (Mesothelae, Liphistiidae), which is constrained to southwestern Japanese islands. It is known for its low vagility and inclination towards population structuring at small to moderate geographical scales (Xu et al., 2015a, 2015b, 2019). Prior to this study, there was taxonomic uncertainty in this genus. Twenty nominal *Heptathela* species are endemic to Kyushu and the northern and central Ryukyus (Tanikawa and Miyashita, 2014; World Spider Catalog, 2020; Xu et al., 2019), including 19 morphological species and one putative cryptic species. Four species are found in Kyushu, five in Amami, and 11 in Okinawa. Like other liphistiids, species of *Heptathela* lack clear diagnostic characters, are morphologically homogeneous among species, exhibit considerable intraspecific variation in female genital morphology, and/or are rare in collections (missing sexes or adult forms) (Xu et al., 2015c, 2017). As demonstrated for other liphistiid genera, *Heptathela* is potentially well suited to the application of an integrative approach to species delimitation.

In this study, we explore and validate species boundaries in *Heptathela* using molecular data via two steps. First, we analyse mitochondrial *COI* sequence data from 180 specimens of *Heptathela* to identify the lineages that are consistently delimited by a range of species-delimitation methods. Second, for any lineages that are not consistently delimited by these methods, we add mitochondrial and nuclear markers and apply multilocus, coalescent-based methods for species delimitation. The results of our analyses allow us to investigate the taxonomy of the genus using a molecular approach that efficiently identifies and investigates problematic lineages.

## 2. Materials and methods

### 2.1 Taxon sampling and DNA sequencing

We carried out three extensive collection trips to Japanese islands in December 2012, September 2013, and May 2014. Although our sampling was not exhaustive, we attempted to sample specimens from across the entire known range of *Heptathela*, from Kyushu to the central Ryukyu archipelago, based on the type locality information of each recorded *Heptathela* species (Fig. 1). We also searched previously unexplored areas. We made a great effort to apply the same strategy to each sampling location, but we often found no spiders because some species are very rare or have patchy distributions. Furthermore, some locations are urbanized habitats while others are inaccessible because of the lack of roads or the presence of military camps. Nevertheless, eight species were sampled from more than one population (3–14 individuals per population), and 12 species were collected from only one population with 2–10 individuals per species. Further details about sampling can be found in our taxonomic study (Xu et al., 2019).

**Fig. 1.**
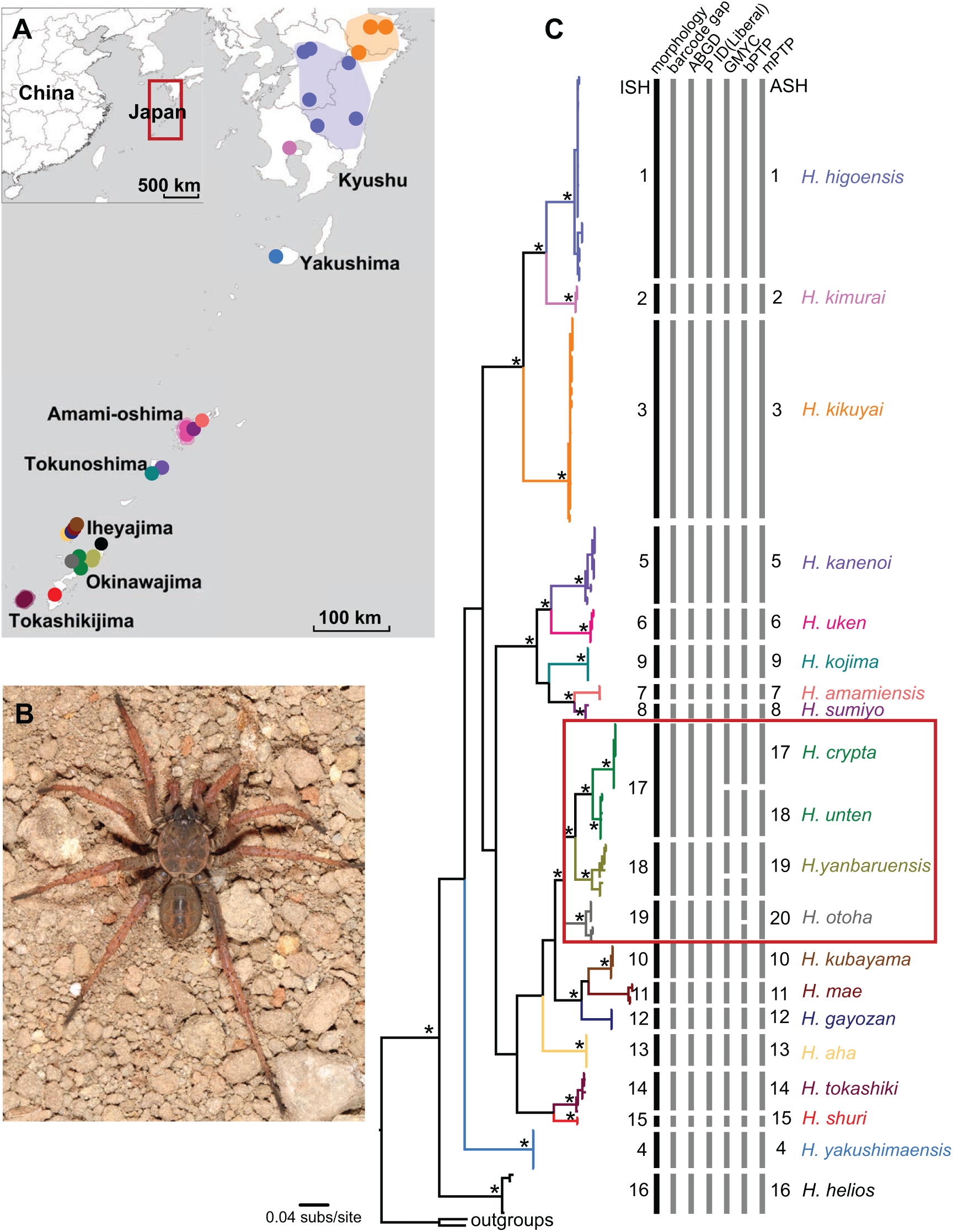
(A) Map showing all sampling localities for species of *Heptathela* on Kyushu and the Ryukyu archipelago (for a more detailed map, see Xu et al., 2019). (B) *Heptathela kojima*, male (body length: 10.40 mm), from Amami-oshima, showing the general somatic morphology. (C) Bayesian tree for 180 samples of *Heptathela*, rooted using two outgroup taxa, *Ryuthela nishihirai* and *R. shimojanai*. Asterisks indicate strongly supported nodes (posterior probability > 0.95, likelihood bootstrap support > 70%). Clades are colour coded to match the map. ISH: initial species hypothesis (19 species), based on morphology and geographic information. ASH: alternative species hypothesis. Vertical bars indicate species delimitations based on morphology and on six different single-locus species delimitation methods. The red box indicates the lineages for which there was disagreement among the six species-delimitation methods.

We selected 180 specimens of *Heptathela* to form the ingroup for our analyses (Supplementary Table S1). In order to root the tree, we included two outgroup taxa from the liphistiid genus *Ryuthela*: *R. nishihirai* (Haupt, 1979) and *R. shimojanai* Xu, Liu, Ono, Chen, Kuntner, & Li, 2017. These taxa are also endemic to Japanese islands (Xu et al., 2017).

From leg muscles of specimens preserved in absolute ethanol, we extracted genomic DNA using the Animal Genomic DNA Isolation Kit (Dingguo, Beijing, China). For all 182 samples, we amplified and sequenced the mitochondrial gene encoding cytochrome oxidase subunit 1 (*COI*) using the primer pair LCO1490/HCO2198 (Folmer et al., 1994). For 35 samples selected from the lineages of *Heptathela* that were not consistently resolved by different single-locus species-delimitation methods (see sections 2.2 and 3.1), we obtained sequence data from one additional mitochondrial gene (16S rRNA, *16S*) and three nuclear markers (28S rRNA, *28S*; internal transcribed spacer 2, *ITS2*; and histone 3, *H3*). DNA from these markers was amplified using the primer pairs 16Sar/16Sbr (Huber et al., 1993), 28S-O/28S-C (Hedin and Maddison, 2001), ITS-5.8S/ITS-28S (White et al., 1990), and H3aF/H3aR (Colgan et al., 1998), respectively. We followed previously reported standard PCR protocols for all genes (Xu et al., 2015a), then manually edited and aligned sequence data in Geneious Prime 2020 (https://www.geneious.com). Owing to a lack of variation in *28S* for the subset of 35 samples, we excluded this gene from our subsequent species-delimitation analyses.

### 2.2. Single-locus species delimitation

We inferred the evolutionary relationships among the sampled individuals of *Heptathela* using the *COI* data set. Our *COI* data set, comprising 676 bp sequences from 180 individuals of *Heptathela*, had 248 variable sites. For each of the three codon sites in the *COI* alignment, the GTR+G substitution model was selected using the Akaike information criterion in jModeltest v2.1.10 (Darriba et al. 2012). We chose not to consider models that allowed a proportion of invariable sites, owing to the shallow evolutionary timescale of *Heptathela* (Jia et al., 2014; Xu et al., 2016).

We performed Bayesian phylogenetic analyses of the *COI* alignment, partitioned by codon site, in MrBayes v3.2.1 (Ronquist et al., 2012). Posterior distributions were estimated by Markov chain Monte Carlo (MCMC) sampling. Samples were drawn every 2000 steps over a total of 20 million steps, with the initial 25% of samples discarded as burn-in. We performed the MCMC analysis in duplicate to check for convergence, and used the *sump* command to check for sufficient sampling. We also carried out a maximum-likelihood phylogenetic analysis of the *COI* alignment, partitioned by codon site, in IQ-TREE v1.6.12 (Nguyen et al., 2015). Node support was estimated using ultrafast bootstrapping with 1000 replicates (Hoang et al., 2018) and the Shimodaira-Hasegawa approximate likelihood-ratio test (SH-aLRT) with 1000 replicates (Guindon et al., 2010). The best-fitting substitution model for each of the three codon sites was selected using ModelFinder in IQ-TREE (Kalyaanamoorthy et al., 2017).

Our initial 19-species hypothesis for *Heptathela* is based on our accompanying morphological study of the genus (Xu et al., 2019). We tested this initial hypothesis by analysing mitochondrial *COI* sequence data with six commonly used single-locus species-delimitation methods. The first two methods were based on genetic distances: DNA barcoding gap (Barrett and Hebert, 2005) and automatic barcode gap discovery (ABGD; Puillandre et al., 2012). The remaining four methods were based on the inferred tree: generalized mixed Yule-coalescent (GMYC; Pons et al., 2006), the P ID(Liberal) method from the Species Delimitation plugin for Geneious (Masters et al., 2011), Bayesian Poisson tree processes (bPTP; Zhang et al., 2013), and multi-rate Poisson tree processes (mPTP; Kapli et al., 2017).

DNA barcoding gap is the most commonly used single-locus technique for species delimitation (Barrett and Hebert, 2005; Hamilton et al., 2014). It uses threshold values to differentiate between inter- and intraspecific divergences. We used MEGA X (Kumar et al., 2018) to calculate pairwise K2P and *p*-distances, as well as mean intra- and interspecific K2P and *p*-distances for each putative species. Unlike DNA barcoding gap analysis, ABGD does not require the sampled individuals to be assigned to putative species (Puillandre et al., 2012). ABGD detects the barcode gap based on the user-defined boundaries for intraspecific variability, then sorts the sampled individuals into candidate species with *p*-values. We performed ABGD analyses online (bioinfo.mnhn.fr/abi/public/abgd/), based on the K2P model, with a minimum intraspecific variability Pmin=0.001, maximum intraspecific variability Pmax=0.1, and minimum gap width X=1.5.

We then used four tree-based methods for species delimitation. The P ID(Liberal) method, implemented in the Species Delimitation plugin in Geneious, tests different species delimitations that are defined *a priori* by the user (Masters et al., 2011). We used the phylogeny inferred in our Bayesian analysis as the guide tree to calculate the mean probability of the ratio of intra- to interspecific genetic distances for the initial 19-species hypothesis for *Heptathela*.

The bPTP method, an updated version of the original PTP with Bayesian support values, assumes independent exponential distributions to model the branch lengths for speciation and for coalescence (Zhang et al., 2013). The newly developed mPTP, on the other hand, fits an independent exponential distribution to the branch lengths of each species (Kapli et al., 2017). We implemented bPTP on an online server (species.h-its.org/ptp/), using the maximum-likelihood phylogeny as the input tree. We ran the bPTP analysis for 500,000 steps, with a thinning of 500 and burn-in proportion of 0.1. We performed species delimitations using likelihood and MCMC analyses on the maximum-likelihood tree in mPTP v0.2.4 (Kapli et al., 2017). In both analyses, we allowed differences in rates of coalescence among species and specified a minimum branch length of 0.0001. We ran MCMC analyses for 100 million steps, sampling every 10,000, and discarded the first 10% of samples as burn-in. Two independent MCMC analyses were run to check for convergence. Analyses starting from the maximum-likelihood species delimitation, random delimitation, and null delimitation all gave the same result.

The GMYC method uses likelihood to identify species boundaries by detecting the transition point between the speciation process and intraspecific lineage coalescence (Pons et al., 2006). We used the single-threshold model in the “splits” package (Ezard et al., 2009) for R v3.6.3 (R Core Team, 2020). The ultrametric guide tree was inferred using Bayesian analysis in BEAST v1.8.4 (Drummond et al., 2012). We compared two clock models (strict clock and relaxed lognormal clock; Drummond et al. 2006) and two tree priors (birth-death model with and without incomplete sampling) using marginal likelihoods estimated by stepping-stone sampling (Xie et al., 2011) with 100 path steps. The birth-death model provides a suitable tree prior for data sets with a mixture of intra- and interspecies sampling (Ritchie et al., 2017). To estimate the divergence times, we used a previous estimate of the *COI* substitution rate from other spider lineages (Bidegaray-Batista et al., 2014) to set a normal prior (mean 0.015, standard deviation 0.001) on the substitution rate. The posterior distribution was estimated by MCMC sampling, with samples drawn every 5000 steps over a total of 50 million steps. We performed the analyses in duplicate to check for convergence and used Tracer to check for sufficient sampling.

### 2.3. Multilocus species delimitation

To investigate the lineages that were subject to inconsistent species delimitations across the six single-locus analyses described above (Fig. 1, red box), we performed multilocus species delimitation using two mitochondrial and two nuclear markers from 35 samples of *Heptathela*. The data set consisted of sequences from mitochondrial *COI* (*n*=35, length 681 bp, 98 variable sites), mitochondrial *16S* (*n*=34, length 491 bp, 46 variable sites), nuclear *H3* (*n*=34, length 360 bp, 17 variable sites), and nuclear *ITS2* (*n*=34, length 344 bp, 3 variable sites). The best-fitting substitution models were HKY+G (*16S* and *COI*), F81 (*ITS2*), and SYM+G (*H3*). As above, we chose not to consider models that allowed a proportion of invariable sites. We concatenated the two mitochondrial markers and treated them as a single locus. We then concatenated the two nuclear markers because they had small numbers of variable sites.

We used two methods, Bayesian Phylogenetics and Phylogeography (BPP; Yang, 2015) and Bayes-factor delimitation (Grummer et al., 2014), to compare the four competing species hypotheses generated from our analyses of the *COI* data (see section 3.1). BPP has been shown to be effective in using information from gene trees to identify the boundaries between closely related species (Flouri et al., 2018; Yang and Rannala, 2017). Unguided Bayesian species delimitation was conducted in BPP v4.14 (Flouri et al., 2018) to explore different species-delimitation models and changes in the topology of the species tree (A11 analysis).

For the priors on population sizes (*θ*) and divergence times (*τ*), we used inverse-Gamma priors with *α* = 3. The *β* parameter was adjusted according to the mean estimate of nucleotide diversity for *θ* (*β* = 0.2) and node height for *τ* (*β* = 0.08). We performed the analyses using the reversible-jump MCMC algorithm 0 (*ε* = 2), and algorithm 1 (*α* = 2, *m* = 1), each run in duplicate to check for convergence. Analyses were run for 100,000 MCMC steps, with samples drawn every 5 steps and with the first 20% of samples discarded as burn-in.

Bayes-factor delimitation compares different species-tree models using their marginal likelihoods. We performed Bayes-factor delimitation in *BEAST (Heled and Drummond, 2010) using a strict molecular clock, a birth-death model for the species-tree prior, and piecewise-linear population sizes. Analyses were run for 50 million MCMC steps, with samples drawn every 5,000 steps and with the first 10% of samples treated as burn-in. We ran the analyses in duplicate to check for convergence and used Tracer to check for sufficient sampling. For each of the four competing species models, we estimated the marginal likelihood using stepping-stone sampling. Bayes factors (BF) were interpreted following the recommendations of Grummer et al. (2014), with 2*ln*BF > 10 being considered as decisive support for a hypothesis. Where a comparison between two models was inconclusive, we favoured the simpler model.

## 3. Results

### 3.1. Single-locus species delimitation

Bayesian and maximum-likelihood methods inferred similar phylogenetic trees (Fig. 1, Supplementary Fig. S1). The initial species hypothesis (i.e., 19 species) was supported by three of the six species-delimitation methods that we compared: barcoding gap analysis, ABGD, and P ID(Liberal) (Fig. 1; Supplementary Tables S2 and S3). We detected a distinct gap between intra- and interspecific genetic distances for 19 hypothetical species of *Heptathela*, ranging from 4.2% to 5.7% for K2P distances and from 4.0% to 5.4% for *p*-distances (Fig. 2). The smallest mean interspecific and largest mean intraspecific distances were 6.0%/5.7% and 2.1%/2.0% (K2P/*p*-distance), respectively.

**Fig. 2.**
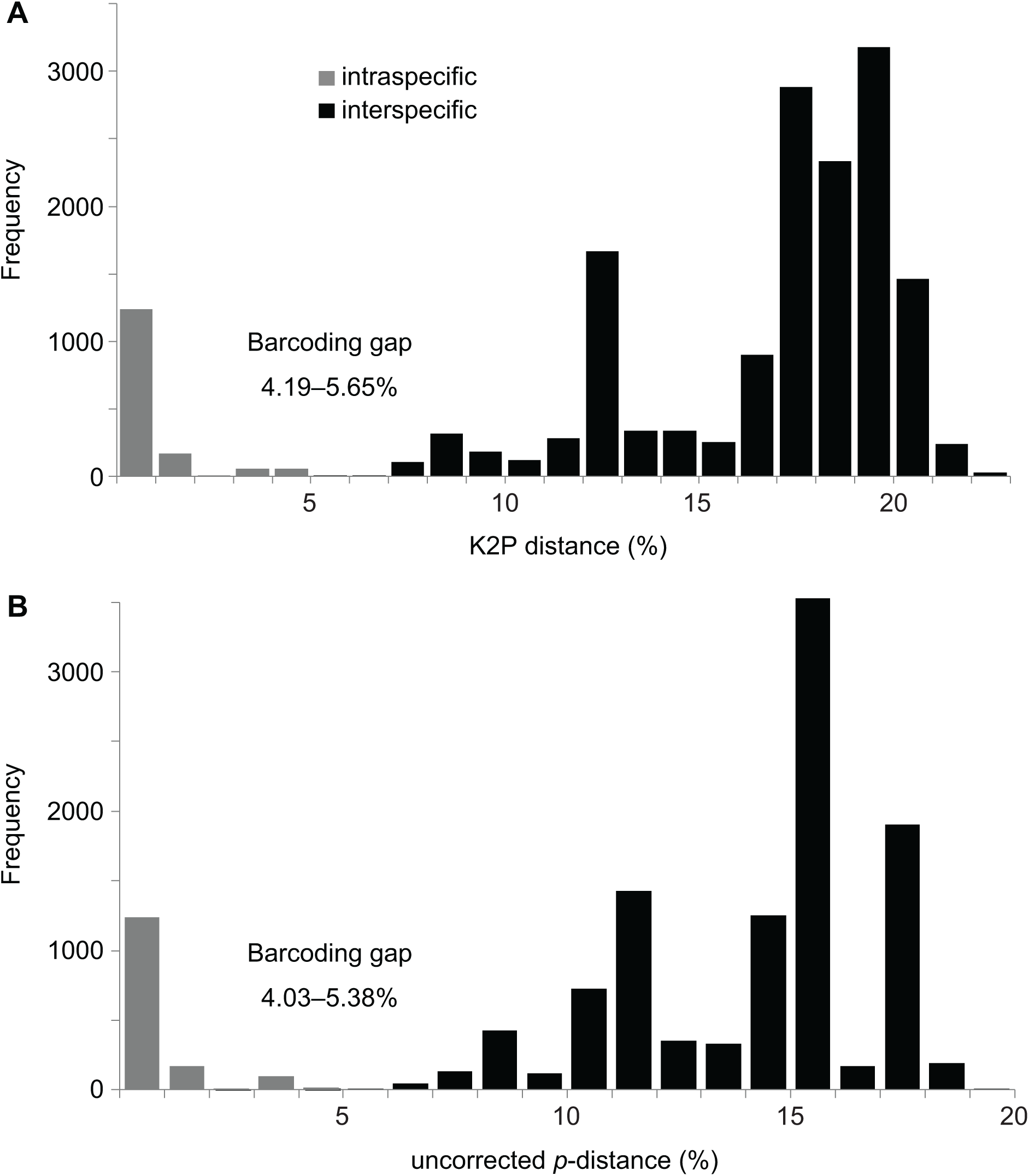
DNA barcoding gap for COI sequences from 19 putative species of *Heptathela*, based on (A) K2P distances and (B) *p*-distances.

The initial species hypothesis was not supported by our analyses using GMYC, bPTP, and mPTP, which delimited 21, 22, and 20 species, respectively. Nevertheless, 16 putative species were recognized by all six of the methods that we applied to the *COI* data set, and these are supported by morphological diagnosis in our taxonomic study (Xu et al., 2019). Based on these results, we derived four competing species hypotheses that we then tested using analyses of multilocus data (Fig. 1).

### 3.2. Multilocus species delimitation

Our BPP analysis of the multilocus data set yielded strong support for three or four species (posterior probability 1) or five species (posterior probability 0.97), with weaker support for six species (posterior probability 0.56) (Table 1). Bayes-factor delimitation showed that marginal likelihoods estimated using stepping-stone sampling provided the highest support for a model with six species, but found that this was not better than the five-species model (2*ln*BF = 1.08) and only marginally better than the four-species model (2*ln*BF = 5.97) (Table 2). In this case, we favour the four-species model because it is the simplest model that is not worse than the best-supported model. Taken together, our analyses using BPP and Bayes-factor delimitation support the four-species model.

**Table 1.** Posterior probabilities from Bayesian analyses of multilocus data in BPP analyses, computed under four competing species hypotheses for the genus *Heptathela*

| Hypothesis | Delimited species | Posterior probability |  |  |  |
| --- | --- | --- | --- | --- | --- |
|  |  | rjMCMC (0, 2) |  | rjMCMC (1, 2, 1) |  |
|  |  | run 1 | run2 | run 1 | run2 |
| 3 species |  | 1.00 | 1.00 | 1.00 | 1.00 |
|  | *otoha* | 1.00 | 1.00 | 1.00 | 1.00 |
|  | *crypta*+*unten* | 1.00 | 1.00 | 1.00 | 1.00 |
|  | *yanbarunesis* | 1.00 | 1.00 | 1.00 | 1.00 |
| 4 species |  | 1.00 | 1.00 | 1.00 | 1.00 |
|  | *otoha* | 1.00 | 1.00 | 1.00 | 1.00 |
|  | *crypta* | 1.00 | 1.00 | 1.00 | 1.00 |
|  | *unten* | 1.00 | 1.00 | 1.00 | 1.00 |
|  | *yanbarunesis* | 1.00 | 1.00 | 1.00 | 1.00 |
| 5 species |  | 0.97 | 0.97 | 0.97 | 0.97 |
|  | *otoha* | 0.98 | 0.97 | 0.98 | 0.98 |
|  | *crypta* | 1.00 | 1.00 | 1.00 | 1.00 |
|  | *unten* | 1.00 | 1.00 | 1.00 | 1.00 |
|  | *yanbarunesis* 1 | 0.97 | 0.97 | 0.97 | 0.97 |
|  | *yanbarunesis* 2 | 0.99 | 0.99 | 0.99 | 0.99 |
| 6 species |  | 0.55 | 0.55 | 0.56 | 0.55 |
|  | *otoha* 1 | 0.64 | 0.63 | 0.64 | 0.63 |
|  | *otoha* 2 | 0.58 | 0.57 | 0.58 | 0.57 |
|  | *crypta* | 0.99 | 1.00 | 1.00 | 1.00 |
|  | *unten* | 0.99 | 0.99 | 0.99 | 1.00 |
|  | *yanbarunesis* 1 | 0.87 | 0.88 | 0.87 | 0.88 |
|  | *yanbarunesis* 2 | 0.99 | 0.99 | 0.99 | 0.99 |

**Table 2.** Marginal likelihoods of four competing species hypotheses for the genus *Heptathela*, computed from a multilocus data set. The log marginal likelihoods are ranked from highest to lowest; the Bayes factor (BF) is calculated using 2 × (marginal likelihood of species model 1 – marginal likelihood of species model 2), with 2*ln*BF > 10 being considered as decisive support for species model 1.

| **Species model** | **Log marginal likelihood** | **Rank** | **$2\ln\text{BF}$** |
| --- | --- | --- | --- |
| 3 species | -4180.17 | 4 | 40.25 |
| 4 species | -4163.03 | 3 | 5.97 |
| 5 species | -4160.59 | 2 | 1.08 |
| 6 species | -4160.05 | 1 |  |

## 4. Discussion

Our two-step approach to molecular species delimitation has yielded support for 20 putative species in the liphistiid spider genus *Heptathela*. The approach uses fast, single-locus delimitation methods to analyse an extensive taxon sample before applying more intensive multilocus methods to a chosen subset of the taxa. Our results confirm the taxonomic utility of the *COI* barcoding region by showing that a range of species-delimitation methods yield largely congruent estimates of species boundaries. Nevertheless, our approach also demonstrates the advantages of comparing the inferences from multiple methods for species delimitation.

Single-locus species-delimitation methods are routinely used to study taxa that are well differentiated genetically but not morphologically, such as liphistiid and mygalomorph spiders (Satler et al., 2013; Xu et al., 2015a, 2017). However, with the widespread acknowledgement of potential discordance between gene trees and the species tree due to incomplete lineage sorting (e.g., Harrington and Near, 2012; McGuire et al., 2007), species delimitation based on a single locus alone can be misleading. Therefore, multilocus methods based on the multispecies coalescent are expected to have a number of advantages. However, the field has not arrived at any single method that would be preferred for its reliability, cost-effectiveness, and robustness.

Comparison of the performance of the six single-locus methods for species delimitation used in this study is not straightforward. Our previous studies on species limits in other liphistiid genera largely relied on the outcomes of DNA barcoding gap, ABGD, and P ID(Liberal) methods (Xu et al., 2015c, 2017). In the present study, all three of these methods support the initial, morphology-based hypothesis of 19 species. On the other hand, bPTP, mPTP, and GMYC support larger numbers of species. As in previous studies of simulated and empirical data (Luo et al., 2018; Tang et al., 2014), our results suggest that bPTP has a slight tendency to oversplit species. The GMYC method is also prone to delimiting larger numbers of species (Hamilton et al., 2014; Talavera et al., 2013).

An important challenge to species delimitation is the overestimation of species diversity, which can be caused by molecular approaches delimiting populations with geographic or genetic structure (Jackson et al., 2017; Sukumaran and Knowles, 2017). Furthermore, oversplitting can become worse when the amount of sequence data increases (Leaché et al., 2019). Bayes-factor delimitation is also prone to species oversplitting, especially if the same data are used to construct and test the species hypotheses (Grummer et al., 2014; Leaché et al., 2014). This problem of oversplitting is potentially heightened in short-range endemic taxa, such as liphistiid spiders, which often have high levels of geographic or population genetic structure (Chambers and Hillis, 2020; Hedin et al., 2015). However, our analyses of *Heptathela* show that both BPP and Bayes-factor delimitation support one of the more conservative species hypotheses that we considered.

Our species delimitations might have been negatively affected by the limitations on sampling. Many species in *Heptathela* are rare and have small geographical ranges, presenting considerable challenges to fieldwork and sample collection. Incomplete or geographically biased sampling can complicate molecular species delimitation (e.g., Ahrens et al., 2016; Carstens et al., 2013; Linck et al., 2019). For example, if all individuals from each species are collected from a single location, then intraspecific diversity will be underestimated, and barcode gaps will be overestimated. In this study, although 11 of the 16 species delimited using *COI* data were collected from a single population, they are consistent with morphological evidence (Xu et al., 2019). Barcode gaps are probably overestimated and will decrease with further sampling for each of these species. Nevertheless, our delimited species are likely to reflect the underlying evolutionary units in the genus *Heptathela*, based on current sampling.

One can still argue that since the conflicting lineages are in a few locations where the geographic scale of sampling seems more ‘fine grained’ (i.e., multiple populations were sampled within a small geographic area), perhaps the species hypotheses generated from our analyses of *COI* sequence data would be less clear if this denser geographic sampling were applied throughout the genus *Heptathela*. Nevertheless, additional genetic data and increased geographic sampling of multiple populations will allow further testing of these species hypotheses.

In species of *Heptathela* from Kyushu and the Ryukyu archipelago, as in other liphistiid genera, females exhibit considerable intraspecific variation. In contrast, males are recognised with stable morphological characters, but they are very rarely collected (Haupt, 2003; Xu et al., 2015c). During our three extensive collection trips, we found both males and females for 12 species, but we found only females for the other eight species (Xu et al., 2019). For the reasons given above, a purely morphology-based taxonomic revision would be likely underestimate the true species diversity. Therefore, molecular species delimitation offers useful diagnostic evidence for the integrative taxonomy of liphistiids.

Eight *Heptathela* species were collected from more than one population, including two species (*H. higoensis* and *H. kikuyai*) that have wide distributions in Kyushu, and six species (*H. kanenoi, H. uken, H. yanbaruensis, H. otoha, H. tokashiki*, and *H. crypta*) with narrow distributions in the Ryukyu archipelago. Five of these eight species were identified by all six single-locus species-delimitation approaches, whereas the other three species were inconsistently delimited by the six methods. Nevertheless, these species delimitations were confirmed by the multilocus, coalescent-based methods. The remaining 12 species that were each collected from only a single population were also supported by single-locus methods, except *H. unten* was validated by multilocus species-delimitation approaches. The combined evidence from the geographical distributions and molecular species delimitation of *Heptathela* suggests that species diversity has not been overestimated. The Ryukyu archipelago has a higher diversity of species of *Heptathela* than Kyushu, despite the latter having a larger area.

In conclusion, our study has formalized an efficient two-step approach to molecular species delimitation by identifying problematic taxa and applying additional markers and analyses in a focused manner. Our analysis of primitively segmented trapdoor spiders of the genus *Heptathela* confirms the effectiveness of our approach, which can provide reliable species delineation. However, our results also demonstrate the need to evaluate multiple lines of evidence for an objective, repeatable test of species boundaries.

## Supporting information

Figure S1, Tables S1-S3, and nucleotide alignment.

## Acknowledgements

We thank Arong Luo and Chengmin Shi for advice on data analyses. This study was supported by the grants from the National Science Foundation of China (NSFC-31601850; NSFC-31272324), the Hunan Provincial Natural Science Foundation of China (2017JJ3202), a bilateral China and Slovenia exchange project No. 12-8 from China, Bi-CN/18-20-022 from Slovenia, the Singapore Ministry of Education AcRF Tier 1 grant (R-154-000-A52-114), the Slovenian Research Agency (P1-0255, P1-10236, J1-672), and a visiting scholarship from China Scholarship Council to X.X. In part, analyses relied on the resources of the National Supercomputing Centre, Singapore (https://www.nscc.sg).

## Appendix Supplementary material

DNA sequences are deposited in GenBank. DNA alignments and Tables S1–S3 as supplementary materials can be found online.

