## Supplementary material for "Molecular species delimitation in the primitively segmented spider genus *Heptathela* endemic to Japanese islands": Figure S1, Tables S1-S3, and nucleotide alignment.: TableS2-3.docx

**Table S2**. Results of the ABGD analyses.

| substitution model | Pmin/Pmax | X | Partition | Prior intraspecific divergence (P) | | | | | | |  |  |
| --- | --- | --- | --- | --- | --- | --- | --- | --- | --- | --- | --- | --- |
|  |  |  |  | 0.001 | 0.0017 | 0.0028 | 0.0046 | 0.0077 | 0.0129 | 0.0215 | 0.0359 | 0.0599 |
| K2P | 0.001/0.1 | 1.5 | Initial | 19 | 19 | 19 | 19 | 19 | 19 | 19 | 19 | 19 |
|  |  |  | Recursive | 54 | 33 | 33 | 25 | 22 | 22 | 20 | 20 | 19 |

**Table S3**. Summary of the mean intraspecific and closest interspecific genetic distance, the mean probability with 95% confidence interval and the intra/inter ratio for 19 initial species hypothesis.

| Putative species | Intraspecific K2P/*p*-distances | Closest P ID(Liberal) species | Closest interspecific K2P/p-distances | P ID(Liberal) | Intra/Inter |
| --- | --- | --- | --- | --- | --- |
| sp19 | 0.0089/0.0088 | sp4 | 0.2101/0.1798 | 0.99 (0.93, 1.0) | 0.04 |
| sp4 | 0/0 | sp19 | 0.2101/0.1798 | 1.00 (0.94, 1.0) | 0.01 |
| sp18 | 0/0 | sp17 | 0.0730/0.0687 | 0.97 (0.82, 1.0) | 0.02 |
| sp17 | 0.0062/0.0062 | sp18 | 0.0730/0.0687 | 0.97 (0.90, 1.0) | 0.12 |
| sp11 | 0.0198/0.0192 | sp10 | 0.0830/0.0774 | 0.95 (0.90, 1.0) | 0.22 |
| sp10 | 0.0209/0.0202 | sp11 | 0.0830/0.0774 | 0.97 (0.94, 1.00) | 0.28 |
| sp12 | 0.0086/0.0085 | sp11 | 0.0892/0.0824 | 0.97 (0.90, 1.0) | 0.12 |
| sp14 | 0.0045/0.0045 | sp13 | 0.0939/0.0868 | 0.97 (0.86, 1.0) | 0.07 |
| sp13 | 0.0015/0.0015 | sp15 | 0.0769/0.0720 | 0.98 (0.87, 1.0) | 0.04 |
| sp15 | 0/0 | sp13 | 0.0769/0.0720 | 0.98 (0.87, 1.0) | 0.02 |
| sp16 | 0/0 | sp12 | 0.1241/0.1114 | 0.98 (0.88, 1.0) | 0.01 |
| sp5 | 0.0081/0.0080 | sp6 | 0.1051/0.0970 | 0.99 (0.95, 1.0) | 0.08 |
| sp6 | 0.0018/0.0018 | sp5 | 0.1051/0.0970 | 0.98 (0.87, 1.0) | 0.03 |
| sp8 | 0.0034/0.0034 | sp7 | 0.0596/0.0566 | 0.96 (0.82, 1.0) | 0.08 |
| sp7 | 0.0011/0.0011 | sp8 | 0.0596/0.0566 | 0.98 (0.84, 1.0) | 0.04 |
| sp9 | 0.0011/0.0011 | sp8 | 0.1034/0.0952 | 0.98 (0.87, 1.0) | 0.02 |
| sp3 | 0.0023/0.0023 | sp1 | 0.1224/0.1108 | 1.00 (0.97, 1.0) | 0.03 |
| sp2 | 0.0010/0.0010 | sp1 | 0.0888/0.0824 | 0.98 (0.87, 1.0) | 0.03 |
| sp1 | 0.0045/0.0044 | sp2 | 0.0888/0.0824 | 0.99 (0.96, 1.0) | 0.07 |
